# Assessing the bioenergy potential of grassland biomass from conservation areas in England

**DOI:** 10.1101/506709

**Authors:** Katherine E. French

## Abstract

Bioenergy may be one of the ‘ecosystem services of the future’ for grasslands managed for conservation as the concept of bio-based economies is embraced worldwide. Although the idea of producing biogas and bioethanol from lignocellulosic material is not new, there are currently few regional-level comparisons of the bioenergy potential of high-diversity grasslands that would establish whether this could be a competitive bioenergy feedstock for farmers. Comparing the chemical composition and biogas yields of biomass samples from 13 grasslands in England and 73 other bioenergy feedstocks reveals that the lignin content of biomass from grasslands managed for conservation was up to 50% less than other bioenergy crops. Grasslands managed for conservation yielded up to 160% more biogas per ton dry matter than cereals or crop waste and only slightly less than Miscanthus. GIS modeling of the estimated biogas yields of grasslands managed for conservation and fields currently sown with Miscanthus show that grasslands are larger (20.57 ha) than Miscanthus fields (5.95 ha) and are projected to produce up to 117% more biogas per average field. Future incorporation of high-diversity grasslands into local and nation-wide energy plans may help reduce global fossil-fuel use in the 21^st^ century.

## 1. Introduction

Global reliance on fossil fuels has led to loss of natural ecosystems and global warming (Butt et al, 2013; Kirschke et al. 2013). In an effort to reduce fossil fuel consumption, producing ethanol from first-generation bioenergy crops like maize and jatropha increased in the 1980s (Openshaw 2000). However, expanded cultivation of primary bioenergy crops has led to the loss of local biodiversity, destruction of soil microbial communities, and increased competition between food and fuel production (Prochnow et al. 2009). Primary energy crops like maize and rapeseed also produce high levels of nitrous oxide (N_2_O), a greenhouse gas 296 times more detrimental than the carbon dioxide (CO_2_) released during fossil fuel consumption, due to high nutrient (nitrogen) requirements (Crutzen et al. 2008). In response, over the past two decades research has focused on developing a number of second and third generation bioenergy crops with a lower environmental impact. These initiatives include producing biogas from crop waste, creating new cultivars of specific crops with enhanced sugar or cellulose contents, and using algae to produce biodiesel (Christian et al. 2008; Jones and Mayfield 2012). Generating bioenergy from plants is now a cornerstone of policies to build stronger bioeconomies in the UK, EU and USA (Burns et al. 2016; EC 2006; McCormick and Kautto 2013).

Producing bioenergy from grasslands may also be a viable alternative to first-generation biofuel production and would promote the preservation of native biodiversity and its associated ecosystem services. Globally, grasslands are increasingly converted to arable land or urban development. They are one of the most threatened biomes yet receive the least conservation attention. For example, temperate grasslands, savannahs and shrublands cover 45.8% of Earth’s terrestrial surface yet only 4.6% of this area is under active protection (Hoekstra et al. 2005). Grasslands provide food for pollinators, flood control, and support ecological food webs sustaining rare plants and animals (Fletcher et al. 2011; Holzschuh et al. 2011; Verdade et al. 2015). Using grasslands currently set aside for conservation for bioenergy production would ensure the maintenance of these ecosystem services while also providing an economic benefit to farmers.

Although the idea of producing biogas and bioethanol from lignocellulosic material is not new (Adler et al. 2009; Herrmann et al. 2013; Van Meerbeek et al. 2016), there are currently few regional-level comparisons of the bioenergy potential of high-diversity grasslands that would establish whether this could be a competitive bioenergy feedstock for farmers. A number of factors may inhibit the production of biogas and bioethanol from grassland biomass. For example, plants typical of grasslands (grasses, forbs and herbs) have tough cell walls composed of cellulose, hemicellulose, and lignin. Lignin tightly binds hemicellulose and cellulose together and fermentation (anaerobic digestion) is necessary to break these bonds to produce biogas and/or ethanol. Grassland biomass is often rejected as a suitable bioenergy feedstock due to its lignin content (Frigon and Giuiot 2010; Triolo et al. 2012). Indeed, a number of international initiatives now focus on breeding crops like barley with lower levels of lignin by using CRISPR/cas9 to induce targeted mutations in cinnamyl alcohol dehydrogenase (CAD), which regulate lignin biosynthesis (Kalluri et al. 2014). However, pre-treating lignocellulosic biomass can increase the biogas yields of substrates with high lignin levels. Steam explosion can separate lignin from hemicellulose and cellulose and can double biogas yields (Hendricks and Zeeman 2009). Fungi, such as *Trichodermo* spp., can also be used to break down lignin before the biomass is added to the digester increasing biogas yields by up to 400% (Muthangya et al. 2009; Wagner et al. 2013). The bacteria used as inoculum in anaerobic digesters can also be optimized to break down lignin (Sun et al. 2013). For example, *Clostridium thermocellum, Comamonas testosteroni*, and *Pseudonocardia autotrophica* contain endoglucanases, exoglucanases, xylanases, and lignolitic enzymes highly effective in degrading plant cell walls (Himmel et al. 2007; Liao et al. 2016).

Despite increased interest as grasslands as a source of bioenergy, the bioenergy output of grasslands compared to other current bioenergy feedstocks is unclear. Here, I estimate the biogas output of three different types of grasslands common to Europe: (1) unimproved grasslands, which are high in biodiversity and offer multiple ecosystem services; (2) restored grasslands, which are former arable fields; and (3) improved grasslands sown with ryegrass (*Lolium perenne* L.), clover (*Trifolium pratense* L., *Trifolium repens* L.) and lucerne (*Medicogo sativa* L.). I specifically chose to assess biogas yield instead of ethanol yield because lignocellulosic feedstocks are more suitable for biogas production. In addition, anaerobic digesters in England (and more broadly, Europe) currently use lignocellulosic materials (e.g. crop waste) to produce biogas and electricity, not ethanol. I then compared the lignocellulosic composition and biogas outputs of these grasslands to 73 other bioenergy feedstocks. Using Oxfordshire, England as a case study, I then conducted a regional analysis of the potential biogas yield of grasslands managed for conservation versus fields sown with Miscanthus. I specifically chose to estimate the potential biogas yields of a single county due to previous objections that the potential land available for bioenergy production is overestimated at the national level (Russelle et al. 2007; Steubing et al. 2010). The present study focuses primarily on the suitability of biomass from grasslands managed for conservation as a bioenergy feedstock and the potential energy yields of these agricultural landscapes. Excluded from this analysis are the economic costs and benefits of bioenergy production from grasslands.

## 2. Methods

### 2.1 Study area

The county of Oxfordshire is located in south-east England. The county has a maritime temperate climate with an annual rainfall of between 570 −750 mm depending on elevation (Killick *et al*. 1998). The primary crops are wheat (*Triticum aestivum* L.), barley (*Hordeum vulgare* L.), and rapeseed (*Brassica napus* L.). Miscanthus and Short Rotation Coppice are currently grown as bioenergy crops on a small scale. There are currently six anaerobic digesters in the county that process lignocellulosic biomass (crop waste, cereals, maize, and ryegrass) (The National Non-Food Crops Center (NNFCC) database, http://www.nnfcc.co.uk/) but none use grassland biomass as a feedstock.

### 2.2 Site selection and vegetation surveys

In July 2015, biomass samples were collected from 13 grasslands. These sites consisted of seven unimproved meadows, four restored meadows, and two improved grasslands. All samples were collected from working farms in Oxfordshire, England to reflect real agricultural conditions. This is particularly important, as most studies on bioenergy output from grasslands are based on biomass samples from experimental plots which may not reflect the species composition of real fields. To determine the species-composition and richness for each field vegetation surveys were conducted at each site. To ensure comparability between fields, I designated a 10 m x 10 m area for survey and forage collection at each site. These sample areas were not selected beforehand because the area sampled was based on the farmer’s decision on the day of the site visit. The presence of grazing livestock, fertilizer application, specific conservation regulations, and farmer interest in the forage quality of specific fields influenced farmer choice. To determine species composition, five 1m^2^ quadrats were randomly placed within each field and the species present in each quadrat were recorded. Abundance was determined as the number of quadrats each species occurred in. Plants were identified using Fitter et al. (1984).

### 2.3 Biomass sample collection and analysis

At each site, biomass samples of ca. 150 grams were collected. Biomass sample collection protocols were adapted from guidelines used for hay-bale sampling developed by the National Forage Testing Association (NFTA) (http://foragetesting.org/) and consultation with forage experts from the Agri-Food and Biosciences Institute (AFBI) (Belfast, Northern Ireland) (http://www.afbini.gov.uk/). To ensure comparability among samples, grasslands were sampled at the same time of day (10 am). While walking in a zig-zag pattern in each field, handfuls of grass were cut ca. 10 cm from the ground with shears at ca. 20 different locations. To ensure an accurate representation of the vegetation composition of the field, all plant species collected in the process of sampling were included in the sample. During collection, grass was placed in a canvas bag to limit changes in forage sugar composition due to increased heat and bacterial activity. Samples were kept at room temperature. Samples were oven dried at 60°C and milled on the same day of collection. Wet chemistry was used to establish Dry Matter (DM) content, sugar, fiber, protein, and lignin content of each sample. Sugar content (water soluble carbohydrate, WSC) was determined by modifying the method created by McDonald and Henderson (1964). Crude protein (CP) was determined using the Kjeldahl method (Association of Official Analytical Chemists 1990). Neutral detergent fiber (NDF) and acid detergent fiber (ADF) were determined using Refluxing method (Van Soest et al. 1991). All sample analyses were performed at the Agri-Food and Biosciences Institute agricultural research center in Hillsborough, England.

### 2.4 Collection of comparative data on bioenergy feedstocks

To compare the lignocellulosic composition of species-rich grass to other bioenergy feedstocks, data on 73 contemporary bioenergy crops from two databases, *Phyllis2* (https://www.ecn.nl/phyllis2/) and *Feedipedia* (http://www.feedipedia.org/), were collected (**Table 1**). Only samples with data on cellulose, hemicellulose and lignin content were included in the analysis. An attempt was made to include at least three examples of each feedstock but this was not possible for all crops. The biogas yield and methane content of the biomass samples from Oxfordshire were compared to a subset of these bioenergy feedstocks (cereals, crop wastes, grass, Miscanthus, legumes, rapeseed, and switchgrass). The bioenergy yield for five feedstocks (newsprint, agave, bamboo, hemp, and kenaf) was not calculated due to absence of dry matter content (DM) data.

**Table 1.** List of contemporary bioenergy feedstocks that were compared to biomass from species-rich grasslands. The lignocellulosic composition of biomass from species-rich grasslands was compared to all feeds listed in Table 1. The biogas yield of biomass from species-rich grasslands was only compared to those feedstocks marked with a *.

| Feedstock | No. | Species (if applicable) |
| --- | --- | --- |
| **Agave** | 1 | *Agave* L. |
| **Bamboo** | 1 | *Bambuseae* sp. Kunth ex Dumort |
| **\*Cereals** | 12 | wheat (*Triticum aestivum* L.), barley (*Hordeum vulgare* L.), oats (*Avena sativa* L.), and maize (*Zea mays* L.) |
| **\*Crop waste** | 8 | corn stover (maize stalks) and straw from wheat, barley and oats |
| **\*Grass** | 15 | Timothy (*Phleum pratense* L.), orchard grass (*Dactylis glomerata* L.), Bromegrass (*Bromus* sp. Scop), Big Bluestem (*Andropogon gerardi* Vitman), Tall Fescue (*Festuca arundinacea* Schreb.), Reed Canary Grass (*Phalaris arundinacea* L.), and verge grass |
| **\*Hemp** | 2 | *Cannabis sativa* L. |
| **\*Kenaf** | 2 | *Hibiscus cannabinus* L. |
| **\*Legumes** | 12 | lucerne (*Medicago sativum* L.), clover (*Trifolium* spp), and soybean (*Glycine max* (L.) Merr.) |
| **\*Miscanthus** | 3 | *Miscanthus x giganteus* Keng |
| **Newsprint/Paper** | 6 | recycled paper, newsprint, and domestic paper waste |
| **\*Rapeseed** | 3 | *Brassica napus* L. |
| **\*Sisal** | 2 | *Agave sisalana* Perrine |
| **Sugarcane** | 2 | *Saccharum* sp. L. |
| **\*Switchgrass** | 4 | *Panicum virgatum* L. |
| **Total** | **73** |  |

### 2.5 Estimated bioenergy output

Biogas yield was calculated based on the chemical composition of each substrate using the Buswell formula (C_c_H_h_O_o_N_n_S_s_ + {(4c − h − 2o + 3n + 2s)/4} H_2_O → {(4c − h + 2o + 3n + 2s)/8} CO**2** + {(4c + h − 2o − 3n − 2s)/8} CH_4_+ nNH_3_ + sH_2_S) (Symons and Buswell 1933; Teghammer 2013; Triolo et al. 2012). Cellulose, hemicellulose, and lignin content were used to calculate the potential bioenergy yield of each feedstock because these are the main substrates converted to biogas in anaerobic digestion. Protein and fat/lipid content was not available for all samples so they were excluded from the analysis. The protein and fats/lipids are usually low for the lignocellulosic materials analyzed in this study so this should make little difference to the total bioenergy yield. As there are three chemical formulas for lignin (C_9_H_10_O_2_, C_10_**H**_12_O_3_, and C_11_**H**_14_O_4_), the molar mass, carbon yield and methane yield were calculated for each and the average of the three was used. To calculate the carbon and methane yields, I used V = nRT/p, where n = amount of substance (mol), R = gas constant (L atm K^−1^ mol^−1^), T = absolute temperature (K), and p = absolute pressure of the gas (atm). In this analysis, R was set at 0.08205747 L atm K^−1^ mol^−1^, T was set at 273.15 K, and p was set at 1 atm according to previously established protocols for estimating biogas yield (Teghammer 2013; Richards et al. 2001). **Supplementary Table 1** shows the biogas yield from cellulose, hemicellulose and lignin based on the Buswell Formula. The biogas yield of lignin and hemicellulose is similar to fat/lipid (C_57_ H_104_O_6_) (1.4 Normal Meter Cubed (Nm^3^)/kg) although the methane concentration of fats/lipids is much higher (70%). The biogas yields of protein (C_5_H_7_O_2_N) (1.0 Nm^3^/kg) and carbohydrate (C_6_H_12_O_6_) (0.8 Nm^3^/kg) are similar to cellulose but lower than hemicellulose and lignin. However, the methane outputs of protein, carbohydrate, cellulose, hemicellulose and lignin are similar (~50%).

To calculate the biogas output, I calculated the biogas yields of cellulose, hemicellulose and lignin for 1 ton dry matter of each sample based on previous established protocols (see Teghammer 2013; Rittmann et al. 2001). Briefly, this can be summarized in the following equation:

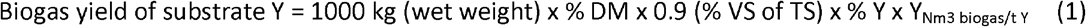

The DM of each sample is used as the total solids (TS) of the sample. To allow for up to 10% of the substrate to be consumed by bacteria during the anaerobic digestion process, the percent volatile solids (VS) of the total solids was set at 0.9. In the equation, Y refers to the % DM of cellulose, hemicellulose and lignin and Y_Nm3 biogas/tY_ refers to the biogas yield of each compound. This is 0.83 for cellulose, 1.2 for hemicellulose, and 1.25 for lignin respectively. The total methane yield (Nm^3^/t) of each sample was calculated based on the methane yields for cellulose (0.50), hemicellulose (0.54) and lignin (0.46). To calculate methane concentrations, I divided total methane content by the total biogas content.

### 2.6 Comparison of grassland and bioenergy crop area and yield

To compare the area covered by grasslands managed for conservation and bioenergy crops, data on the total area of SSSIs, grasslands under agro-environmental schemes, and fields sown with Miscanthus and Short Rotation Coppice in Oxfordshire was obtained from Natural England (http://www.geostore.com/environment-agency/WebStore?xm=environment-agency/xml/ogcDataDownload.xml). This data is public sector information licensed under the Open Government License v1.0. All maps were created using ArcGIS^®^ software by Esri. All records were screened to remove duplicates (e.g. fields associated with more than one scheme) which would inflate actual estimates of grassland coverage. The potential yield of grassland biomass was estimated at 8-10 tons dry matter (tDM) per ha based on previous estimates of average grassland biomass yields in the UK and northern Europe (Amon et al. 2006; Rösch et al. 2009; Seppälä et al. 2009).^1^ The potential biomass yield of Miscanthus was estimated at 10-14 t/ha based on previous research by the Biomass Energy Center and the UK Forestry Commission (Biomass Energy Center 2008). The average yield per ha for grasslands managed under agro-environmental schemes was based on the lowest average biomass yield for grasslands (8 t/ha) and the average biogas yield of species-rich grasslands estimated in section 3.3. The biogas yield per ha for Miscanthus was based on the lowest average biomass yield for Miscanthus (10t/ha) and the average biogas yield of Miscanthus estimated in section 3.3. These predicted yields are based on actual field sizes (ha). These estimates are used for heuristic purposes and actual biomass yields per hectare may vary from field to field and from year to year given variation in species composition (and in the case of Miscanthus, genotype) and annual rainfall (Clifton-Brown et al. 2001).

### 2.7 Statistical Analysis

I used one-way analysis of variance (ANOVA) to determine whether there were any statistically significant differences in lignin, hemicellulose, and cellulose contents of the forage samples from Oxfordshire. To determine which species were associated high levels of lignin, hemicellulose and cellulose content, I used indirect gradient analysis using Redundancy Analysis (RDA) followed by a Monte Carlo permutation test with 499 permutations on log-transformed data. Based on this data, I created General Additive Models (GAMs) for cellulose, hemicellulose, and lignin to determine which species were associated with increased yields of each material. Only species with a response variable of P < 0.05 were included. To determine the relationship between vegetation species-richness and bioenergy yield, correlation analysis (Pearson’s Correlation Coefficient) was used followed by a *t*-test to establish significance of the r values (Crawley 2011). The same test was performed to determine whether cellulose, hemicellulose and lignin were correlated with biogas yield. To compare the lignocellulosic composition of species-rich grass and other bioenergy feedstocks, I used correspondence analysis (CA) on log-transformed data. To determine whether there were any statistically significant differences in lignin, hemicellulose, cellulose content, and bioenergy yield of species-rich grasslands and other bioenergy crops, I used ANOVA. Correlation analysis and ANOVA were performed in R version 3.2.2 (“Fire Safety”) and RDA and CA was performed in Canoco (version 4.5, Lepš and Šmilauer 2003).

## 3. Results

### 3.1 Lignocellulosic composition and biogas yield of grassland biomass

Cellulose (*F*_2,10_ = 0.333, P = 0.725) and lignin (*F*_2,10_ = 2.408, P = 0.14) content did not vary significantly among the three grassland types (**Table 2; Supplementary Materials Table 2**). However, there was a marginally significant difference in hemicellulose content among grassland types (*F*_2,10_ = 3.775, P = 0.06). Unimproved grasslands had the highest average cellulose content while restored grasslands had the highest hemicellulose content. Biogas yields varied significantly among the three grassland types (*F*_2,10_ = 6.243, P = 0.017). Restored grasslands had the highest average biogas yield followed by unimproved grasslands, although there was no significant difference between the two (*t* = 0.699, P = 0.50). The average biogas yield of improved grasslands was 30% lower than that of unimproved grasslands.

**Table 2.** Comparison of the cellulose, hemicellulose, lignin and biogas yields of unimproved, restored and improved grasslands. Data in the table shows the mean ± one standard error.

|  | Cellulose<br>(% DM) | Hemicellulose<br>(% DM) | Lignin<br>(% DM) | Biogas Yield<br>(Nm<sup>3</sup> / ton) |
| --- | --- | --- | --- | --- |
| Unimproved Grassland | 24.1 ± 2.2 | 22.9 ± 1.7 | 10.3 ± 2.2 | 625.1 ± 28.3 |
| Restored Grassland | 22.1 ± 3.8 | 24.7 ± 2.9 | 12.5 ± 3.7 | 657.9 ± 46.9 |
| Improved Grassland | 26.4 ± 2.3 | 14 ± 3.7 | 1.3 ± 4.8 | 437.5 ± 60.1 |

### 3.2 Effect of vegetation species richness on lignocellulosic composition and biogas yield

Vegetation species-richness showed a strong positive correlation with lignin content (r = 0.71, *t* = 3.30, df = 11, P = 0.008) and was not significantly correlated with either hemicellulose content (r = 0.29, *t* = 0.99, df = 11, P = 0.34) or cellulose content (r = 0.46, *t* = −1.73, df = 11, P = 0.11). There was a marginally significant positive correlation between vegetation species and biogas yield (r = 0.50, *t* = 1.92, df =11, P = 0.08) (**Figure 1A**). Hemicellulose (r = 0.94, *t* = 9.46, df = 11, P < 0.001) and lignin (r = 0.66, *t* = 2.877, df = 11, P = 0.01) were strongly positively correlated with biogas yield. There was no correlation between biogas yield and cellulose content (r = −0. 24, *t* = −0.99, df = 11, P = 0.46). RDA analysis showing the relationship between lignocellulosic composition and biogas yield is depicted in **Figure 1 B**. To identify which plants were correlated with increase hemicellulose and lignin content, and thus, potentially greater biogas yields, GAMS were created for hemicellulose and lignin. Yellow oat grass (*F* = 6.07, P = 0.019), nettle (*Urtica dioica* L.) (*F* = 10.72, P = 0.003) and ryegrass (F= 8.19, P = 0.008) were associated with increases in hemicellulose (**Figure 1 C**). Increased lignin levels were associated with increased abundances of orchard grass (*F* = 6.69, P = 0.01), Yorkshire Fog (*Holcus lanatus* L.) (*F* = 9.22, P = 0.005), common reed (*F* = 33.78, P < 0.001), and Floating Sweet Grass (*Glyceria fluitans* (L.) R. Br.) (*F* = 33.78, P < 0.001) (**Figure 1 D**).

**Figure 1.**
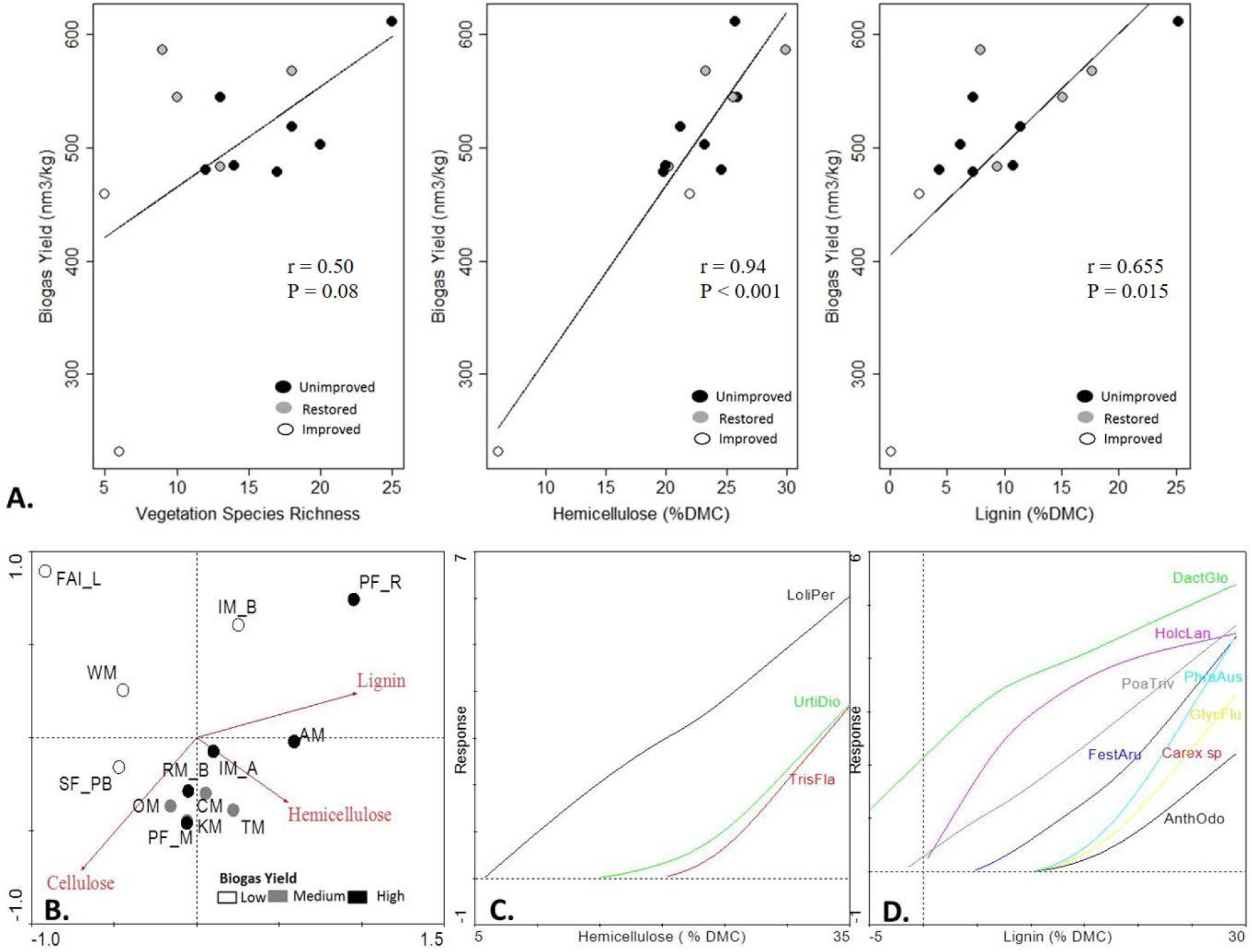
Effect of vegetation species-richness and composition on estimated grassland biogas yields. A. Correlation analysis showing the relationship between species-richness, hemicellulose and lignin content and biogas yield. B. RDA analysis showing lignocellulosic composition of grass samples from Oxfordshire. Samples are coded based on biogas yield (high, medium, low). RDA explains 45% of the variation in species among samples and 100% of the correlation between species and the lignocellulosic components (*F* = 1.957, P = 0.026). C. and D. Generalized Additive Models (GAMs) showing the association of particular species with hemicellulose and lignin. Species are labeled by the first four letters of the genus and first three letters of the species. Only species with P < 0.05 were included in each model.

### 3.3 Comparison to other bioenergy feedstocks

There were significant differences in cellulose (*F*_14,69_ = 6.642, P < 0.001), hemicellulose (*F*_14,69_ = 6.79, P < 0.001), and lignin (*F*_14,69_ = 4.03, P < 0.001) content among crop types among the different feedstocks. Hemp, sisal and kenaf had the highest cellulose content. Cereals, biomass from species-rich grasslands, and legumes had the lowest cellulose content. Switchgrass, Miscanthus, and crop waste had the highest average hemicellulose levels and kenaf, hemp and legumes had the lowest. Bamboo, Miscanthus, and rapeseed had the highest average lignin levels while agave, cereals and hemp had the lowest. Correspondence analysis indicates that the lignocellulosic composition of biomass from species-rich grasslands is most comparable to crop waste, grass, and switchgrass (**Figure 2A**).

**Figure 2.**
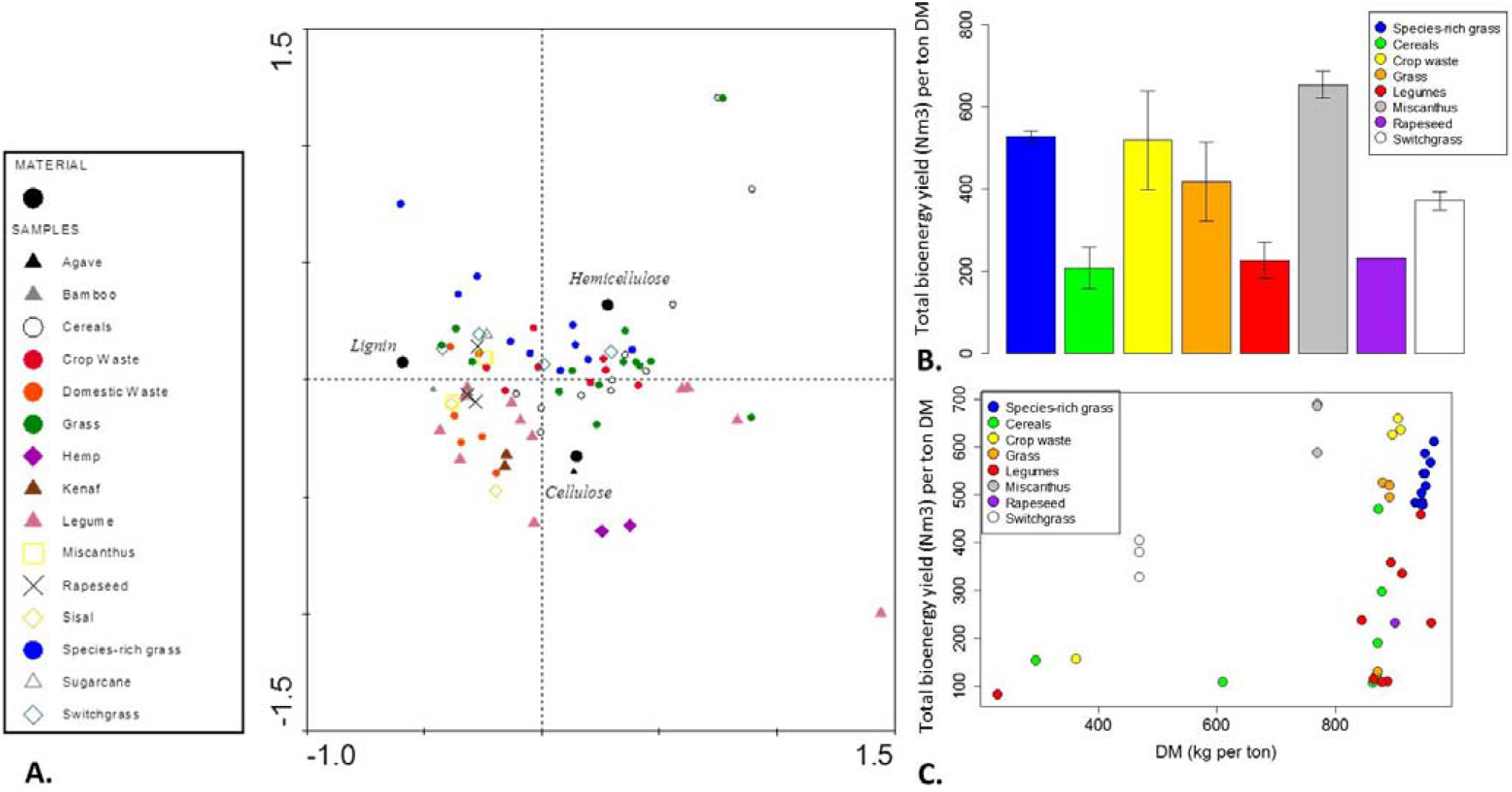
Comparison of grasslands to other bioenergy feedstocks. A. Correspondence analysis showing the lignocellulosic composition of bioenergy feedstocks. B. Estimated biogas yield of bioenergy feedstocks based on 1 ton dry matter. Bar plot shows mean ± standard error. C. Correlation between dry matter yield and biogas yield. In the legends, “species-rich grass” refers to biomass from unimproved and restored grasslands.

Biogas yield varied significantly among bioenergy feedstock types (**F**_7,35_= 5.33, P < 0.001) (**Table 3; Figure 2B; Supplementary Materials Table 3** contains the results for each sample). The average biogas yield of species-rich grass was up to three times higher than that of cereals, legumes, grass, and rapeseed. There was no significant difference between the biogas yield of biomass from species rich grasslands and crop waste (*t* = −1.57, P = 0.12), Switchgrass (*t* = −1.53, P = 0.13), and Miscanthus (*t* = 1.47, P = 0.15). Despite differences in biogas and methane yield, methane concentrations of all feedstocks were similar (around 50%). Biogas yield was positively correlated with dry matter content (r = 0.4, *t* = 2.7, df = 40, p = 0.01) (**Figure 2C**).

**Table 3.** Comparison of lignocellulosic composition and biogas yield of biomass from grasslands managed for conservation to other bioenergy feedstocks. In the table, averages for each feedstock are represented as the mean ± one standard error. Feedstocks marked with ‘nd’ (‘no data’) indicate no DM content was available preventing the calculation of biogas yield.

| Feedstock | Cellulose<br>(% DM) | Hemi-<br>cellulose<br>(%DM) | Lignin<br>(%DM) | DM (kg/t wet<br>weight) | Biogas yield<br>(Nm<sup>3</sup> / t DM) | Methane<br>yield (Nm<sup>3</sup> / t<br>DM) |
| --- | --- | --- | --- | --- | --- | --- |
| **Biomass from**<br>**grasslands**<br>**managed for**<br>**conservation** | 23.4 ± 3.1 | 23.5 ± 1.6 | 11.2 ± 1.8 | 949.9 ± 48.2 | 527.2 ± 39 | 268.5 ± 20 |
| **Agave** | 55.8 ± 10.6 | 15.3 ± 5.5 | 6.8 ± 6.1 | nd | nd | nd |
| **Bamboo** | 39.5 ± 10.6 | 17.6 ± 5.55 | 25.2 ± 6.1 | nd | nd | nd |
| **Cereals** | 24.9 ± 4.2 | 18 ± 2.2 | 4.9 ± 2.4 | 679 ± 104.2 | 207.3 ± 62.6 | 108.6 ± 32.6 |
| **Crop Waste** | 38.2 ± 4.7 | 26.6 ± 2.4 | 9.8 ± 2.7 | 775.6 ± 77.3 | 518.9 ± 75.6 | 266.7 ± 38.7 |
| **Domestic**<br>**Waste** | 38.8 ± 5.1 | 15.8 ± 2.6 | 16 ± 2.7 | nd | nd | nd |
| **Grass** | 20.6 ± 4.0 | 22.9 ± 2.1 | 8.9 ± 2.5 | 880.8 ± 86.3 | 417.7 ± 75.6 | 216 ± 38.7 |
| **Hemp** | 68.5 ± 7.8 | 12.5 ± 4 | 4.5 ± 2.3 | nd | nd | nd |
| **Kenaf** | 53.9 ± 7.8 | 14 ± 4 | 12.1 ± 4.5 | nd | nd | nd |
| **Legumes** | 22.7 ± 4.2 | 10 ± 2.2 | 7.1 ± 4.5 | 829.3 ± 71.9 | 226.3 ± 58.2 | 115.2 ± 29.8 |
| **Miscanthus** | 44.6 ± 6.6 | 25.8 ± 2.2 | 21.2 ± 2.4 | 768 ± 104.2 | 653.9 ± 84.2 | 328.2 ± 42.1 |
| **Rapeseed** | 42 ± 6.6 | 22 ± 3.46 | 19.3 ± 3.8 | 903.5 ± 122.9 | 232.3 ± 135.1 | 116.1 ± 69.3 |
| **Sisal** | 58.9 ± 7.8 | 15.3 ± 3.4 | 17.9 ± 4.5 | nd | nd | nd |
| **Sugar Beet** | 34.8 ± 7.8 | 23 ± 4 | 17.7 ± 4.5 | nd | nd | nd |
| **Switchgrass** | 37.1 ± 5.9 | 29.17 ± 4.1 | 15.7 ± 3.9 | 470 ± 104.2 | 370.8 ± 84.3 | 189 ± 43.2 |

### 3.4 Estimated bioenergy yields of grasslands managed for biodiversity conservation

The area, average field size, and total biomass yield varied significantly among SSSIs, fields managed under agroenvironmental schemes, and Miscanthus (*F*_20,7585_ = 45.49, P < 0.001) (**Figure 3A; Supplementary Materials Table 4**). In Oxfordshire, 107 SSSI grasslands occupy 2,201.53 ha^2^. The average site size is 20.57 ± 0.06 ha^2^. Using the minimum estimate of 8 tDM per ha, the minimum potential biomass yield from these areas would be 17,612.17 tDM while using the maximum estimate of 10 tDM ha gives a total of 22,015.34 tDM. Areas managed under agroenvironmental schemes cover a total of 30,331.13 ha^2^. This consists of 7,479 grasslands managed under 19 different agro-environmental schemes (Entry and High Level Stewardship). Most sites are managed under schemes that maintain permanent grassland under article 13 (3,187 sites) and permanent grasslands with low (1,471 sites) and very low inputs (1,330 sites) not located in Severely Disadvantaged Areas (SDA) or 2 2 above the Moorland Line (ML).^2^ Overall, the average field size is 8.31 ± 0.92 ha and the largest fields are managed to protect the habitat of breeding wetland birds. Using the minimum estimate of 8 tDM per ha, the minimum potential biomass yield from these areas would be 242,649 tDM while using the maximum estimate of 10 tDM ha gives a total of 303,311.3 tDM. In comparison, the average area used for bioenergy crop production under Defra’s bioenergy crops scheme from 2003-2013 was 174.18 ha^2^, with 119.06 ha^2^ planted with Miscanthus. Miscanthus fields were 5.95 ± 1.58 ha^2^ on average. Using the minimum estimate of 10 tDM per ha, the minimum potential biomass yield from these areas would be 1,190.6 tDM while using the maximum estimate of 14 tDM ha gives a total of 1,664.84 tDM.

**Figure 3.**
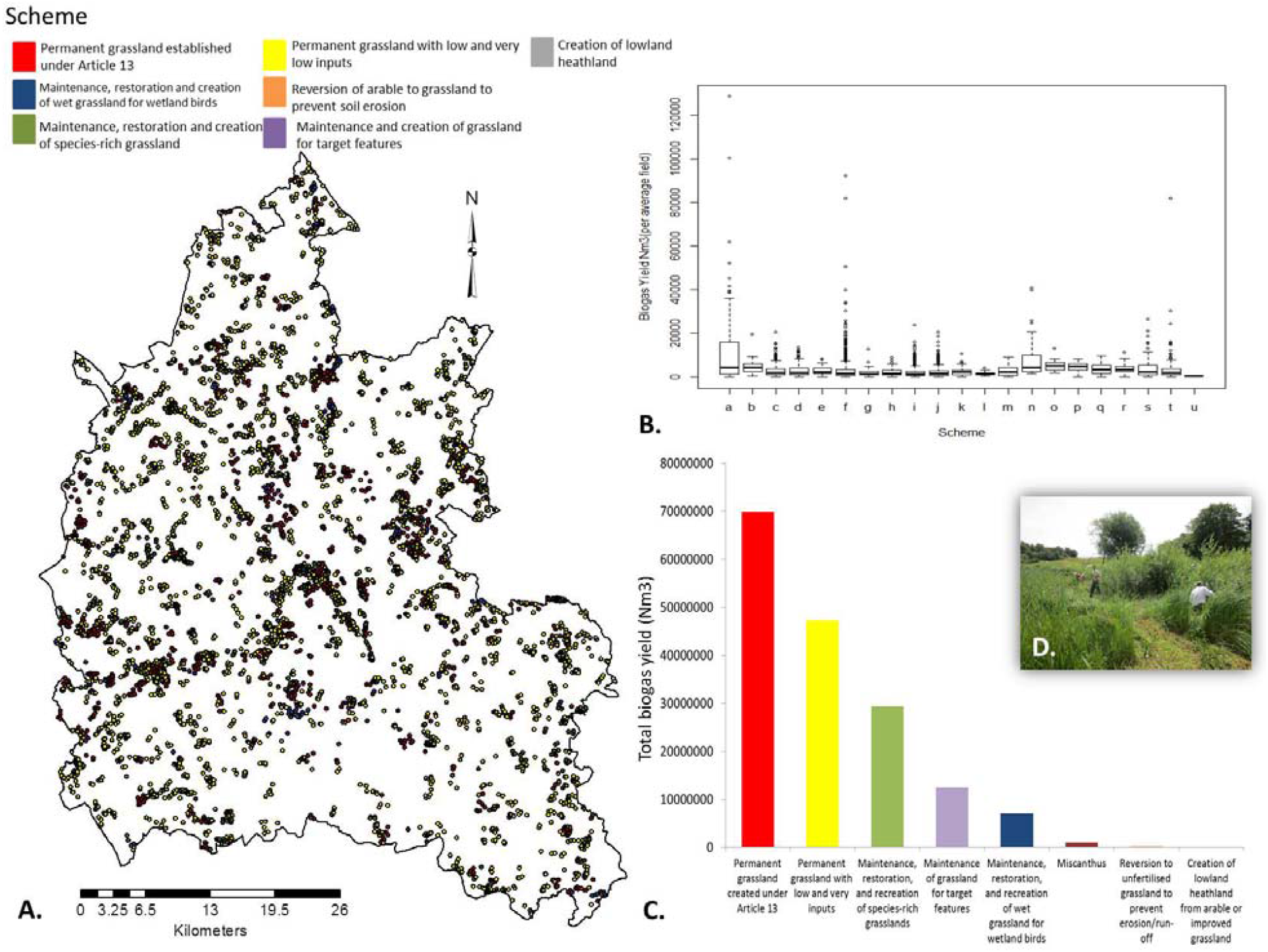
Regional level analysis of grassland bioenergy potential. A. Distribution of grasslands managed under current agro-environmental schemes in Oxfordshire, England. B. Boxplot showing the average biogas yield per field according to scheme. In the diagram, a = SSSIs, b = Miscanthus, f = permanent grassland (unpaid under Article 13), n = maintenance of wet grassland for breeding waders, and o = restoration of wet grassland for breeding waders. The rest of the schemes can be found in Supplementary Materials Table 4. C. Total estimated biogas yield of land managed under each scheme. D. Example grassland managed for conservation in Oxfordshire. Volunteers typically cut and burn the harvested biomass each summer.

The average biogas yield per field varied significantly among SSSIs, Miscanthus, and the fields managed under the 19 different agroenvironmental schemes (*F*_20,17585_ = 45. 54, P < 0.001) (**Figure 3B**). SSSIs had the highest yield (104,867 ± 3,191Nm^3^), followed by wetlands maintained for breeding waders (73,415 ± 6,174 Nm^3^), and Miscanthus (48,283 ± 8,041Nm^3^). Permanent grasslands managed with low and very low inputs (18,723 ± 3,305 Nm^3^ and 15,605 ± 3,317 Nm^3^ respectively) and newly created heathlands had the lowest average yields per field (3,568 ± 33,161 Nm^3^). In order to produce yields competitive to Miscanthus fields producing on average 8,110.72 Nm^3^/ha, grasslands would need to be at least 9.5 ha in size. This would include roughly 43% (46) of grassland SSSIs in the county and 8% (616) of grasslands under agro-environmental schemes. At the scheme level, permanent grasslands maintained under Article 13 (69,879,544 Nm^3^), permanent grasslands with low inputs (outside SDA and ML) (24,598,946 Nm^3^), permanent grasslands with very low inputs (outside SDA and ML) (20,756,514 Nm^3^), SSSIs (11,223,459 Nm^3^), and maintained species-rich, semi-natural grasslands (9,490,812 Nm^3^) had the highest total estimated biogas yield. Miscanthus fields produced 965,209 Nm^3^ in total. The schemes with the lowest potential total biogas yield were unfertilized grasslands created to prevent erosion (184,619 Nm^3^), permanent grasslands managed with low inputs (128,892 Nm^3^) and newly created heathlands (3,379 Nm^3^) (**Figure 3C**).

## 4. Discussion

The suitability of grassland biomass as a bioenergy feedstock is highly debated. Here, I show how vegetation species richness and species composition positively impact grassland biogas yield. I demonstrate that lignocellulosic composition and biogas yield of grasslands managed for conservation are comparable to other bioenergy feedstocks. Finally, my regional estimation of the potential bioenergy yields of grasslands managed for conservation illustrates that they could potentially yield more biogas compared to fields sown with Miscanthus fields due to larger average field sizes and the amount of land currently used for grassland conservation.

### 4.1 Vegetation species-composition affects grassland bioenergy yield

Species-composition plays a strong role in the lignocellulosic composition and potential biogas yields of grasslands. The significantly higher estimated biogas yields of unimproved and conservation grasslands compared to improved grasslands may be due to the presence of tall species with high levels of lignin and hemicellulose. For example, biomass samples with the highest potential biogas yield contained reeds (*P. australis*), Orchard grass (*D. glomerata*), Yellow Oat-Grass (*T. flavescens*), and Giant Fescue (*F. gigantea*). All of these plants are taller than 100 cm. To date, there is little quantitative data on the biogas yields of specific grassland species. However, *P. australis* has been shown to produce high levels of biogas during anaerobic digestion (Lin 2012; Melts et al. 2013, 2014; Prochnow et al. 2009). In the US, bioethanol yields of prairies are also found to be higher on grasslands dominated by tall C_4_ grasses (*Panicum virgatum* L, *Andropogon gerardii* Vitman, and *Sorghastrum nutans* (L.) Nash) than grasslands with a higher species-richness (Adler et al. 2009). These results challenge current research on the effect of biodiversity on the bioenergy yields of grasslands. Increasing species-richness may not directly result in increased bioenergy yields. Rather, selecting the right species may have a greater effect. To balance the needs of conservation while optimizing bioenergy potential, grasslands under restoration could be sown with species amenable to both goals.

### 4.2 Biomass from grasslands managed for conservation is comparable to current bioenergy feedstocks

Despite claims that grassland biomass is high in lignin (Frigon and Giuiot 2010), this study presents the first comparative analysis showing that grasslands managed for conservation produce biomass similar in lignocellulosic composition to other bioenergy feedstocks. This suggests that there should be no technological barrier to producing biogas from grasslands. The estimated biogas yield of biomass from grasslands managed for conservation was also comparable to other feedstocks such as Switchgrass and Miscanthus. All three feedstocks had a high DM content, which may explain these results. Although these results are encouraging, they are based on biomass harvested in July and on theoretical calculation of bioenergy potential based on chemical composition and further research using batch scale digesters is needed to confirm these claims. However, these results do agree with other studies in Europe and the US that report high biogas/ethanol yields from grasslands. For example, Steubing et al. (2010) found that biogas yields from meadows and pastures (17.4 GJ/t DM) were slightly higher than the yield from forest wood (15.8 GJ/t DM), waste paper and cardboard (17 GJ/t DM), and current bioenergy crops (17.3 GJ/t DM). Tilman et al. (2006) also found that low-input high diversity grasslands in the US produced three times more bioenergy than Switchgrass (based on estimating ethanol output at 0.255 L per kg^−1^ dry matter). In order to establish the true bioenergy yield of grasslands managed for conservation, a larger program of research sampling a greater number of grasslands and preparing biomass with currently available pretreatments to break down lignin (e.g. those used for Miscanthus) should be conducted.

### 4.3 Integrating agriculture, biodiversity conservation and bioenergy production

Globally, the area of grassland managed for conservation is increasing rapidly as traditional pastoral systems decline in the UK, Europe, and other parts of the world (Hodgson et al. 2005; Hoekstra et al. 2005). Turning the unused biomass from these areas into bioenergy would give these landscapes a new purpose while also promoting the conservation of these habitats. In Oxfordshire, SSSIs, wetlands maintained for breeding birds, and permanent grasslands cover large areas and had the highest estimated biogas yields by field and in total ha^2^. Similar patterns of grassland cover are seen across England (French, *unpublished data*). In many cases, these grasslands are not used for livestock production (for hay or grazing) and produce excess biomass that is usually burned in the late summer or autumn. Turning this biomass into bioenergy could provide farmers with an additional source of income. For example, in Oxfordshire (and in Europe) permanent grasslands protected under Article 13 cover a large geographic area but farmers do not receive payments for maintaining these landscapes. Farmers could produce electricity and heating by processing grassland biomass in farm-scale anaerobic digesters (which could be used or sold). Grass is an attractive bioenergy crop: farmers are familiar with managing grasslands, do not need specialized machinery to harvest hay, and harvesting grass for bioenergy fits into current arable time cycles. However, if grasslands currently managed for conservation were used for bioenergy production, specific monitoring programs would need to be put in place to ensure local biodiversity (e.g. specific rare species) or landscape quality does not decline. Some habitats (e.g. wetlands maintained for breeding waders) require specific management activities, specifically, late biomass cuts. This should not affect bioenergy yield however, particularly as increased DM content appears to correlate with increased bioenergy yields (Heinsoo et al. 2011).

If bioenergy production from grasslands is carried out on a large scale, it could contribute to national economic growth and reduced greenhouse gas emissions, as seen in Sweden and Germany (Jones and Salter 2013; Parmlind 2014). A lifecycle analysis, taking into the costs associated with harvesting and processing grassland biomass for biogas production, would be useful to both farmers considering biogas production and policy makers. However, the county-level analysis of potential grassland bioenergy yields presented here should be read with caution. First, not all of these grasslands might be suitable for bioenergy production due to terrain (e.g. sloping hills) or prior use (e.g. traditional grazing) (Ciello 2009; Stuebing et al. 2010). To avoid potential land-use conflicts, land traditionally used for grazing could be set aside and protected from any potential enrollment in national bioenergy schemes. Second, the variation in biomass among grasslands is currently unknown. I have based my estimates at 8 tDM per ha which is considered ‘average’ for the region. However, chalk grasslands (which are low in soil nitrogen) produce shorter and less dense biomass than more fertile fen meadows. A citizen science initiative, where farmers and conservation workers measure sward height and biomass, would lead to a more accurate picture of the potential bioenergy yield of grasslands managed for conservation.

## 5. Conclusion

Finding sustainable sources of energy to replace fossil fuels will be one of the greatest challenges of the 21^st^ century. Producing bioenergy from grasslands is a potential practical solution that has been underexplored to date. Biomass from grasslands managed for conservation is comparable in lignocellulosic composition to other bioenergy feedstocks and even contains less lignin than one of the most popular and lauded bioenergy crops, Miscanthus. In addition, the estimated biogas yield of grassland biomass exceeds that of current substrates used for anaerobic digestion. On a regional level, grasslands currently managed for conservation are on average up to four times larger than fields sown with bioenergy crops and occupy dramatically more ha^2^. Giving a ‘new purpose’ to these landscapes could reduce biomass waste, boost farmer interest in conservation, and provide farmers with a new source of income. However, the full potential of grassland biomass as a bioenergy crop can only be realized with changes to current agroenvironmental policies. Current policies could be adapted to increase monetary incentives for farmers to harvest grassland biomass for bioenergy in the form of a cash payment per grassland ha^2^ used for bioenergy production. Policy makers could also help facilitate the use of grasslands for bioenergy production by providing farmers with access to resources (e.g. training in AD technology, different methods for processing biomass for digestion to optimize biogas yield, etc.). By taking these steps, using grasslands for bioenergy production could contribute to reducing reliance on fossil fuels, decreasing cultivation of primary bioenergy crops, and achieving national goals of reducing carbon emissions by 2050, all while conserving native biodiversity.

## Acknowledgements

The author wishes to thank Natural England, FAI Farms, Earth Trust, Berkshire, Buckinghamshire, and Oxfordshire Wildlife Trust (BBOWT) and local farmers and graziers for providing access to their grasslands and L. Turnbull for comments on earlier versions of this paper.

## Footnotes

1 Previous studies report yields from semi-natural grasslands ranging from ca. 3-25 t/ha (DeHaan et al. 2009; Seppälä et al. 2009; Tilman et al. 2006). For example, the yield of semi-natural grassland in the American prairies is 3.7 t/ha while the yield of *Phragmltes australis* dominated wetlands in Sweden is 10 t/ha (Lin 2012). The same is true for Miscanthus, with biomass yields ranging from 8-27 t/ha (Bauen et al. 2010; Christian et al. 2008; Himken et al. 1997; Jørgensen 1997; Kahle et al. 2001). Given the wide variation in biomass yields for grasslands and Miscanthus, I used the more conservative yield estimates for both crops based on UK and European sources.

2 SDA and ML refer to areas where farming is challenging due to rough terrain.

## References

Adler, P.R. et al., 2009. Plant species composition and biofuel yields of conservation grasslands. Ecological Applications, 19(8), pp.2202–2209.

Amon, T. et al., 2006. Methane production through anaerobic digestion of various energy crops grown in sustainable crop rotations. Bioresource Technology, 98(17), pp.3204–3212.

Anon, Miscanthus (*Miscanthus* x *giganteus*) for Biofuel Production - eXtension. Available at: http://www.extension.org/pages/26625/miscanthus-miscanthus-x-giganteus-for-biofuel-production [Accessed October 30, 2015].

Bauen, A.W. et al., 2010. Modelling supply and demand of bioenergy from short rotation coppice and Miscanthus in the UK. Bioresource Technology, 101(21), pp.8132–8143.

Biomass Energy Center (BEC), 2008, Miscanthus. Sources of Biomass. Available at: http://www.biomassenergycentre.org.uk/.

Burns, C., Higson, A. & Hodgson, E., 2016. Five recommendations to kick-start bioeconomy innovation in the UK. Biofuels, Bioproducts And Biorefining, 10(1), pp.12–16.

Butt, N. et al., 2013. Biodiversity Risks from Fossil Fuel Extraction. Science, 342(6157), pp.425–426.

Christian, D.G., Riche, A.B. & Yates, N.E., 2008. Growth, yield and mineral content of *Miscanthus × giganteus* grown as a biofuel for 14 successive harvests. Industrial Crops and Products, 28(3), pp.320–327.

Clifton-Brown, J.C. et al., 2001. Performance of 15 Genotypes at Five Sites in Europe. Agronomy Journal, 93(5), p.1013.

Crawley, M.J., 2011. Statistics: an introduction using R., Hoboken: John Wiley & Sons.

Crutzen, P.J. et al., 2008. N2O release from agro-biofuel production negates global warming reduction by replacing fossil fuels. Atmos. Chem. Phys., 8(2), pp.389–395.

DeHaan, L., S. Weisberg, D. Tilman, and D. Fornara. 2009. Agricultural and biofuel implications of a species diversity experiment with native perennial grassland plants. Agriculture, Ecosystems, and the Environment 137: 33–38.

Energy Research Centre of the Netherlands, Phyllis2, database for biomass and waste. Available at: https://www.ecn.nl/phyllis2.

European Commission (EC), 2005. New perspectives on the knowledge-based bio-economy: conference report, Brussels: European Commission.

Fitter, R., et al., 1984. Collins guide to the grasses, sedges, rushes, and ferns of Britain and Northern Europe, London: Collins.

Fletcher, R.J. et al., 2011. Biodiversity conservation in the era of biofuels: risks and opportunities. Frontiers in Ecology and the Environment, 9(3), pp.161–168.

Frigon, J.-C. & Guiot, S.R., 2010. Biomethane production from starch and lignocellulosic crops: a comparative review. Biofuels, Bioproducts and Biorefining, 4(4), pp.447–458.

Heinsoo, K. et al., 2011. Reed canary grass yield and fuel quality in Estonian farmers’ fields. Biomass and Bioenergy, 35(1), pp.617–625.

Herrmann, C., A. Prochnow, M. Heiermann, and C Idler. 2013. Biomass from landscape management of grassland used for biogas production: Effects of harvest date and silage additives on feedstock quality and methane yield. Grass and Forage Science. 69. 10.1111/gfs.12086.

Himken, M. et al., 1997. Cultivation of Miscanthus under West European conditions: Seasonal changes in dry matter production, nutrient uptake and remobilization. Plant and Soil, 189(1), pp.117–126.

Himmel, M.E. et al., 2007. Biomass recalcitrance: Engineering plants and enzymes for biofuels production. Science, 315(5813), pp.804–807.

Hodgson, J.G. et al., 2005. How much will it cost to save grassland diversity? Biological Conservation, 122(2), pp.263–273.

Hoekstra, J.M. et al., 2005. Confronting a biome crisis: global disparities of habitat loss and protection. Ecology Letters, 8(1), pp.23–29.

Holzschuh, A. et al., 2011. Expansion of mass-flowering crops leads to transient pollinator dilution and reduced wild plant pollination. Proceedings of the Royal Society B-Biological Sciences, 278(1723), pp.3444–3451.

Jones, P. & Salter, A., 2013. Modelling the economics of farm-based anaerobic digestion in a UK whole-farm context. Energy Policy, 62, pp.215–225.

Jones, C.S. & Mayfield, S.P., 2012. Algae biofuels: versatility for the future of bioenergy. Current Opinion in Biotechnology, 23(3), pp.346–351.

Kahle, P. et al., 2001. Cropping of Miscanthus in Central Europe: biomass production and influence on nutrients and soil organic matter. European Journal of Agronomy, 15(3), pp.171–184.

Kalluri, U.C. et al., 2014. Systems and synthetic biology approaches to alter plant cell walls and reduce biomass recalcitrance. Plant Biotechnology Journal, 12(9), pp.1207–1216.

Kirschke, S. et al., 2013. Three decades of global methane sources and sinks. Nature Geoscience, 6(10), pp.813–823.

Lepš, J. & Šmilauer, P., 2003. *Multivariate analysis of ecological data using CANOCO*, Cambridge: Cambridge University Press.

Liao, J.C. et al., 2016. Fuelling the future: microbial engineering for the production of sustainable biofuels. Nature Reviews Microbiology, 14(5), pp.288–304.

Lin, S., 2012. Wetland biomass - Chemical benefits and problems with biogas usage, Available at: http://www.diva-portal.org/smash/record.jsf?pid=diva2%3A535260&dswid=-7907 [Accessed October 30, 2015].

McCormick, K. & Kautto, N., 2013. The Bioeconomy in Europe: An Overview. Sustainability, 5(6), pp.2589–2608.

Melts, I. et al., 2013. Comparison of two different bioenergy production options from late harvested biomass of Estonian semi-natural grasslands. Energy, 61, pp.6–12.

Melts, I., Heinsoo, K. & Ivask, M., 2014. Herbage production and chemical characteristics for bioenergy production by plant functional groups from semi-natural grasslands. Biomass and Bioenergy, 67, pp.160–166.

Muthangya, M., Mshandete, A.M. & Kivaisi, A.K., 2009. Two-Stage Fungal Pre-Treatment for Improved Biogas Production from Sisal Leaf Decortication Residues. International Journal of Molecular Sciences, 10(11), pp.4805–4815.

Openshaw, K., 2000. A review of Jatropha curcas: an oil plant of unfulfilled promise. Biomass & Bioenergy, 19(1), pp.1–15.

Parmlind, E., 2014. Energy analysis of farm-based biogas plants in Sweden. Uppsala: SLU, Dept. of Energy and Technology. Available at: http://stud.epsilon.slu.se/6765/.

Prochnow, A. et al., 2009. Bioenergy from permanent grassland – A review: 1. Biogas. Bioresource Technology, 100(21), pp.4931–4944.

Richards, B.K. et al., 2001. Methods for kinetic analysis of methane fermentation in high solids biomass digesters. Biomass and Bioenergy, 1(2), pp.65–73.

Rittmann, B.E., 2001. Environmental biotechnology: principles and applications, Boston: McGraw-Hill.

Rösch, C. et al., 2009. Energy production from grassland – Assessing the sustainability of different process chains under German conditions. Biomass and Bioenergy, 33(4), pp.689–700.

Russelle, M.P. et al., 2007. Comment on “Carbon-negative biofuels from low-input high-diversity grassland biomass”. Science (New York, N.Y.), 316(5831), p.1567; author reply 1567.

Seppälä, M. et al., 2009. Biogas production from boreal herbaceous grasses – Specific methane yield and methane yield per hectare. Bioresource Technology, 100(12), pp.2952–2958.

Steubing, B. et al., 2010. Bioenergy in Switzerland: Assessing the domestic sustainable biomass potential. Renewable and Sustainable Energy Reviews, 14(8), pp.2256–2265.

Sun, L., Müller, B. & Schnürer, A., 2013. Biogas production from wheat straw: community structure of cellulose-degrading bacteria. Energy, Sustainability and Society, 3(1), p.15.

Teghammer, A., 2013. Biogas Production from Lignocelluloses: Pretreatment, Substrate Characterization, Co-digestion, and Economic Evaluation. PhD thesis. Göteborg, Sweden: Doktorsavhandlingar vid Chalmers Tekniska Högskola.

Tilman, D., Hill, J. & Lehman, C., 2006. Carbon-Negative Biofuels from Low-Input High-Diversity Grassland Biomass. Science, 314(5805), pp.1598–1600.

Triolo, J.M. et al., 2012. Biochemical methane potential and anaerobic biodegradability of non-herbaceous and herbaceous phytomass in biogas production. Bioresource Technology, 125, pp.226–232.

Van Meerbeek, K., S. Ottoy, M. de Andres Garcia, B. Muys, and M. Hermy. 2016. The bioenergy potential of Natura 2000 – a synergy between climate change mitigation and biodiversity protection. Frontiers in Ecology and the Environment, https://doi.org/10.1002/fee.1425

Verdade, L.M., Piña, C.I. & Rosalino, L.M., 2015. Biofuels and biodiversity: Challenges and opportunities. Environmental Development, 15, pp.64–78.

Wagner, A.O., Schwarzenauer, T. & Illmer, P., 2013. Improvement of methane generation capacity by aerobic pre-treatment of organic waste with a cellulolytic *Trichoderma viride* culture. Journal of Environmental Management, 129, pp.357–360.

