## Supplementary Information for "Assessing the bioenergy potential of grassland biomass from conservation areas in England"

**Supplementary Materials**

Table 1 Biogas yield of cellulose, hemicellulose and lignin (per kg) based on the Buswell Formula.

|  | **Cellulose (C_6_H_10_O_5_)** | **Hemicellulose (C_31_H_34_O_11_)** | **Lignin (Average)** |
| --- | --- | --- | --- |
| **H_2_O** | 1 | 17 | 5.5 |
| **CO_2_** | 3 | 14 | 4.25 |
| **CH_4_** | 3 | 17 | 3.67 |
| **Molar mass (g/mol)** | 162.14 | 582.59 | 180.20 |
| **Carbon dioxide volume**  **(V = nRT/p)** | 67.24 | 313.79 | 95.25 |
| **Carbon yield (Nm^3^/ kg)** | 0.41 | 0.53 | 0.53 |
| **Methane volume**  **(V = nRT/p)** | 67.24 | 381.03 | 82.18 |
| **Methane yield (Nm^3^/ kg)** | 0.41 | 0.65 | 0.72 |
| **Total Biogas Yield (per kg)** | 0.82 | 1.19 | 1.25 |
| **Ratio CO_2_ toCH_4_** | 3:3 | 14:17 | 4.24:3.67 |
| **% that is methane** | 50 | 55 | 46 |

Table 2 Vegetation species richness and lignocellulosic content of biomass samples from Oxfordshire. The lignocellulosic composition is in the form % DM.

| Site^[1]^ | Type | Species Richness | Cellulose | Hemicellulose | Lignin |
| --- | --- | --- | --- | --- | --- |
| WM | Unimproved | 12 | 26.03 | 24.58 | 4.3 |
| PF_M | Unimproved | 17 | 28 | 19.78 | 7.3 |
| TM | Unimproved | 18 | 25 | 21.20 | 11.4 |
| IM_A | Unimproved | 13 | 28.43 | 25.86 | 7.3 |
| IM_B | Unimproved | 14 | 23.38 | 19.96 | 10.8 |
| OM | Unimproved | 20 | 28.44 | 23.14 | 6.2 |
| CM | Restored | 13 | 25.69 | 20.25 | 9.4 |
| RM_B | Restored | 9 | 27.39 | 29.86 | 7.9 |
| KM | Restored | 10 | 16.89 | 25.51 | 15.1 |
| AM | Restored | 18 | 18.8 | 23.26 | 17.7 |
| FAI_L | Improved | 6 | 23.39 | 6.08 | 0.1 |
| SF_PB | Improved | 5 | 29.44 | 21.93 | 2.6 |
| PF_R | Unimproved | 25 | 9.45 | 25.75 | 25.2 |

Table 3 Potential biogas yield, methane yield and methane concentration for bioenergy feedstocks based on lignocellulosic composition. Samples from grasslands managed for conservation are in bold and were sampled as part of this study.

| Sample | DMC (%) | DMC (Kg) | VS (at 90%) | Cellulose (%DMC) | Hemicellulose (%DMC) | Lignin (%DMC) | Total biogas yield | Total methane yield | Methane concentration |
| --- | --- | --- | --- | --- | --- | --- | --- | --- | --- |
| Mscnth1 | 0.77 | 769.00 | 692.10 | 0.45 | 0.30 | 0.21 | 689.33 | 347.36 | 0.50 |
| Mscnth2 | 0.77 | 769.00 | 692.10 | 0.45 | 0.30 | 0.21 | 684.29 | 344.71 | 0.50 |
| HordStw | 0.91 | 906.00 | 815.40 | 0.42 | 0.31 | 0.07 | 659.14 | 339.13 | 0.51 |
| TritStw | 0.91 | 910.00 | 819.00 | 0.43 | 0.28 | 0.07 | 634.92 | 325.32 | 0.51 |
| AvnStw | 0.90 | 896.00 | 806.40 | 0.38 | 0.31 | 0.07 | 624.72 | 321.85 | 0.52 |
| Mscnth3 | 0.77 | 769.00 | 692.10 | 0.44 | 0.18 | 0.22 | 588.03 | 292.45 | 0.50 |
| RM_B | **0.95** | **951.40** | **856.26** | **0.27** | **0.30** | **0.08** | **586.03** | **301.91** | **0.52** |
| PF_R | **0.97** | **966.50** | **869.85** | **0.09** | **0.26** | **0.25** | **611.01** | **305.30** | **0.50** |
| AM | **0.96** | **960.00** | **864.00** | **0.19** | **0.23** | **0.18** | **567.14** | **285.57** | **0.50** |
| IM_A | **0.95** | **948.90** | **854.01** | **0.28** | **0.26** | **0.07** | **544.46** | **279.72** | **0.51** |
| KM | **0.95** | **951.70** | **856.53** | **0.17** | **0.26** | **0.15** | **543.95** | **275.99** | **0.51** |
| PhlHy | 0.88 | 880.00 | 792.00 | 0.33 | 0.28 | 0.04 | 525.43 | 271.46 | 0.52 |
| DactHy | 0.89 | 891.00 | 801.90 | 0.32 | 0.27 | 0.05 | 519.83 | 268.58 | 0.52 |
| TM | **0.95** | **951.80** | **856.62** | **0.25** | **0.21** | **0.11** | **517.74** | **262.70** | **0.51** |
| AvnHy | 0.89 | 892.00 | 802.80 | 0.34 | 0.24 | 0.04 | 495.38 | 255.10 | 0.51 |
| OM | **0.94** | **944.90** | **850.41** | **0.28** | **0.23** | **0.06** | **502.79** | **258.20** | **0.51** |
| PF_M | **0.95** | **948.00** | **853.20** | **0.28** | **0.20** | **0.07** | **478.65** | **244.31** | **0.51** |
| IM_B | **0.95** | **946.40** | **851.76** | **0.23** | **0.20** | **0.11** | **484.29** | **245.71** | **0.51** |
| CM | **0.94** | **935.20** | **841.68** | **0.26** | **0.20** | **0.09** | **482.89** | **245.67** | **0.51** |
| WM | **0.94** | **944.00** | **849.60** | **0.26** | **0.25** | **0.04** | **479.82** | **248.11** | **0.52** |
| SF_PB | 0.94 | 944.00 | 849.60 | 0.29 | 0.22 | 0.03 | 458.79 | 237.24 | 0.52 |
| DrdMze | 0.87 | 873.00 | 785.70 | 0.22 | 0.33 | 0.02 | 470.65 | 247.10 | 0.53 |
| SwtchG3 | 0.47 | 470.00 | 423.00 | 0.37 | 0.32 | 0.22 | 404.51 | 204.11 | 0.50 |
| SwtchG4 | 0.47 | 470.00 | 423.00 | 0.45 | 0.31 | 0.12 | 380.48 | 194.08 | 0.51 |
| LucHy | 0.89 | 894.00 | 804.60 | 0.26 | 0.11 | 0.08 | 358.80 | 180.75 | 0.50 |
| LtsHy | 0.91 | 912.00 | 820.80 | 0.24 | 0.07 | 0.10 | 334.96 | 166.38 | 0.50 |
| SwtchG2 | 0.47 | 470.00 | 423.00 | 0.37 | 0.33 | 0.06 | 327.27 | 168.96 | 0.52 |
| AvnCr | 0.88 | 879.00 | 791.10 | 0.14 | 0.19 | 0.03 | 297.90 | 155.29 | 0.52 |
| FAI_L | 0.96 | 961.80 | 865.62 | 0.23 | 0.06 | 0.00 | 232.29 | 118.63 | 0.51 |
| PisSil | 0.84 | 844.00 | 759.60 | 0.19 | 0.09 | 0.04 | 237.71 | 120.43 | 0.51 |
| Rpsd | 0.90 | 901.00 | 810.90 | 0.11 | 0.08 | 0.08 | 232.26 | 116.08 | 0.50 |
| LinM | 0.91 | 906.00 | 815.40 | 0.09 | 0.10 | 0.06 | 223.61 | 113.23 | 0.51 |
| HordCr | 0.87 | 871.00 | 783.90 | 0.05 | 0.15 | 0.01 | 189.19 | 99.92 | 0.53 |
| HordSIl | 0.36 | 363.00 | 326.70 | 0.25 | 0.18 | 0.04 | 156.87 | 80.54 | 0.51 |
| Wht | 0.30 | 295.00 | 265.50 | 0.27 | 0.25 | 0.05 | 153.48 | 79.25 | 0.52 |
| Trtcle | 0.87 | 871.00 | 783.90 | 0.03 | 0.11 | 0.01 | 130.23 | 68.78 | 0.53 |
| TritCr | 0.87 | 870.00 | 783.00 | 0.03 | 0.10 | 0.01 | 123.79 | 65.34 | 0.53 |
| Pisum | 0.87 | 865.00 | 778.50 | 0.07 | 0.07 | 0.00 | 113.80 | 59.44 | 0.52 |
| Soyb | 0.89 | 888.00 | 799.20 | 0.07 | 0.06 | 0.01 | 109.84 | 56.55 | 0.51 |
| SoyM | 0.88 | 879.00 | 791.10 | 0.08 | 0.05 | 0.01 | 108.42 | 55.94 | 0.52 |
| MzSil | 0.61 | 610.00 | 549.00 | 0.08 | 0.09 | 0.02 | 108.98 | 56.42 | 0.52 |
| Mze | 0.86 | 863.00 | 776.70 | 0.02 | 0.09 | 0.01 | 107.04 | 56.72 | 0.53 |
| LtsF | 0.23 | 231.00 | 207.90 | 0.18 | 0.10 | 0.10 | 82.50 | 41.23 | 0.50 |
| SgrBt | 0.16 | 163.00 | 146.70 | 0.06 | 0.06 | 0.01 | 20.18 | 10.45 | 0.52 |

**Table 4 Average biogas yield per hectare of grasslands under current agro-environmental schemes, SSSIs and Miscanthus.** The average biogas yield per field for grasslands under current agro-environmental schemes and SSSIs was based on average biogas yield for species-rich grasslands from Oxfordshire (527.2 Nm^3^) producing 8 tons dry matter per hectare. The average yield per field of Miscanthus was based on the average yield of Miscanthus (653.9 Nm^3^) producing 10 tons dry matter per hectare. For all sites, the actual size (in hectares) of each site was used to determine average biogas yield. The average for each field type was then calculated. The total biomass yield and total bioenergy yield for each scheme was calculated as the sum of all the individual field values within each scheme. All data on the number of farms in each agro-environmental scheme and the size of each parcel of land was obtained from Natural England (see Methods, section 2.6).

|  | Scheme | No. Fields | Avg. Area (ha) | Avg. Yield per Field (Nm^3^) | Total Biomass Yield (tDM/ ha) | Total Biogas Yield |
| --- | --- | --- | --- | --- | --- | --- |
| a | SSSIs | 107 | 20.58 ± 0.63 | 104,867 ± 3,191 | 17,616.48 | 11,223,459 |
| b | Miscanthus | 20 | 5.95 ±1.58 | 48,283 ± 8,041 | 1,190 | 965,209 |
| c | Maintenance of species-rich, semi-natural grassland | 408 | 4.56 ± 0.7 | 23,264 ± 3,585 | 14,896.9 | 9,490,812 |
| d | Restoration of species-rich, semi-natural grassland | 320 | 4.67± 0.72 | 23,787 ± 3,585 | 11,947.52 | 7,611,765 |
| e | Creation of species-rich, semi-natural grassland | 44 | 4.61 ± 1.16 | 23,516 ± 5,911 | 1,624.13 | 1,034,732 |
| f | Permanent grassland (created under Article 13) | 3187 | 4.30 ± 0.64 | 21,924 ± 3,244 | 109,683.79 | 69,879,544 |
| g | Permanent grassland with very low inputs (outside SDA&ML) (organic) | 72 | 3.34 ± 0.99 | 17,034 ± 5,031 | 1,924.99 | 1,226,412 |
| h | Permanent grassland with low inputs (outside SDA & ML)(organic) | 91 | 3.79 ± 0.92 | 19,296 ± 4,707 | 2,756.21 | 1,755,980 |
| i | Permanent grassland with very low inputs (outside SDA & ML) | 1330 | 3.06± 0.65 | 15,605 ± 3,317 | 32,579.68 | 20,756,514 |
| j | Permanent grassland with low inputs (outside SDA & ML) | 1471 | 3.28 ± 0.65 | 18,723 ± 3,305 | 38,610.81 | 24,598,946 |
| k | Permanent grassland with very low inputs | 22 | 4.63 ± 1.51 | 23,614 ± 7,727 | 815.41 | 519,496 |
| l | Permanent grassland with low inputs | 9 | 2.97 ± 2.25 | 15,121 ± 11,456 | 213.62 | 136,100 |
| m | Reversion to unfertilized grassland to prevent erosion/run-off | 8 | 4.78 ± 2.37 | 24,369 ± 12,098 | 305.98 | 194,942 |
| n | Maintenance of wet grassland for breeding waders | 39 | 14.4 ± 1.21 | 73,415 ­± 6,174 | 4,492.8 | 2,862,363 |
| o | Restoration of wet grassland for breeding waders | 34 | 8.83 ± 1.27 | 44,994 ± 6,498 | 2,401.22 | 1,529,815 |
| p | Creation of wet grassland for breeding waders | 13 | 7.14 ± 1.89 | 36,391 ± 9,695 | 742.56 | 473,085 |
| q | Maintenance of wet grassland for wintering waders and wildfowl | 42 | 6.13 ± 1.18 | 31,234 ± 6,010 | 2,059.01 | 1,311,794 |
| r | Restoration of wet grassland for wintering waders and wildfowl | 27 | 6.18 ± 1.39 | 31,489 ± 7,109 | 1,334.45 | 850,177 |
| s | Creation of grassland for target features | 54 | 7.56 ± 1.08 | 38, 513 ± 5, 510 | 3,264.19 | 2,079,617 |
| t | Maintenance of grassland for target features | 307 | 5.29 ± 0.73 | 26, 978 ± 3, 706 | 12,999.61 | 8,282,050 |
| u | Creation of lowland heathland from arable or improved grassland | 1 | 0.7 ± 6.5 | 3, 568 ± 33, 161 | 5.6 | 3,568 |

1. These are field identifiers, used to protect the anonymity of the study participants.
